# Protection against *Neisseria meningitidis* nasopharyngeal colonization relies on antibody opsonization and phagocytosis by neutrophils

**DOI:** 10.1101/2024.09.11.612551

**Authors:** Elissa G. Currie, Olga Rojas, Isaac S. Lee, Khashayar Khaleghi, Alberto Martin, Jennifer Gommerman, Scott D. Gray-Owen

## Abstract

*Neisseria meningitidis* is a human-restricted pathogen that can cause a rapidly progressing invasive meningococcal disease, yet it is also a regular inhabitant of the human nasopharynx. Vaccines that target *N. meningitidis* aim to prevent invasive disease, but their ability to interfere with nasal colonization could effectively eradicate this bacteria in a population, and so is an important target for meningococcal vaccine design. While protection against invasive meningococcal disease is classically attributed to IgG-dependent complement activation and bacterial killing, there remains no indication of what confers protection against nasopharyngeal colonization, making it impossible to deliberately target this stage during vaccine development. Moreover, without understanding what confers protection in this tissue site, it is impossible to understand the level of susceptibility within a population. To address this, we have taken advantage of the CEACAM1-humanized mouse model to characterize immune effectors that protect against nasal carriage of *N. meningitidis*. Protection against nasal colonization could be induced by live mucosal infection or by parenteral immunization with heat-killed bacteria. Mice possessing genetic deficiencies in B cells were used to evaluate the role of B cells and a specific antibody response, while neutrophil and complement depletion were used to evaluate their respective contributions to immunization-induced protection against meningococcal nasal carriage. Despite the essential role for complement killing in preventing invasive meningococcal disease, complement was not required for protection against nasal colonization. Instead, *N. meningitidis-*specific antibodies and neutrophils were both required to protect mice against the nasal infection. Combined, these data suggest that phagocytic bacterial killing is necessary for protection against mucosal colonization by *N. meningitidis*, indicating that nasal immunoglobulin with the ability to promote opsonophagocytosis must be considered as a correlate of protection against meningococcal carriage.

**AUTHOR’S SUMMARY:** *Neisseria meningitidis* can cause devastating and often fatal systemic infections including sepsis and meningitis, yet it frequently lives in the throat of healthy individuals. Vaccines developed against some meningococcal strains allow the individual to resist becoming colonized by the bacteria, an effect that protects them from disease and prevents them from spreading the bacteria to others, while other vaccines effectively protect against disease but still allow the individual to carry the bacteria in their throat. The reason for this difference has remained difficult to explain. Here, we use a ‘humanized’ mouse model that allows *N. meningitidis* infection in the nasal passages to establish that effective protection against nasal colonization requires that antibodies present within the infected mucosal tissues can coat the bacteria so that they are engulfed by neutrophils, a potent bacteria-killing white blood cell that is recruited to the site of infection. These findings suggest that antibodies with the ability to promote neutrophil recognition and killing of *N. meningitidis* should be the goal of future vaccines, and the presence of these can be used to consider an individual’s resistance against this terrible pathogen.

## INTRODUCTION

*Neisseria meningitidis* (*Nme*) is a regular colonizer of the human nasopharynx, estimated to be present in approximately 10% of the general population (1). While nasal colonization is not associated with any clinical manifestations, *Nme* can penetrate the mucosa to cause very severe invasive disease, most commonly manifesting as sepsis and meningitis (2). Due to the rapid progression of disease once onset, meningococcal vaccines are required to reduce the devastating burden of invasive disease. Vaccines targeting *Nme* are approved based on their ability to elicit antibodies that activate complement-mediated bacterial killing, measured by the serum bactericidal assay (SBA), a well-accepted correlate of protection against invasive disease (3).

Vaccines that use *Nme* capsular polysaccharides conjugated to a protein carrier have been very effective at targeting disease caused by serogroups A, C, W and Y (4, 5). These conjugate vaccines reduce nasal colonization and transmission of the pathogen in addition to preventing invasive disease in the vaccinated individual, thereby offering the potential for herd immunity (6, 7). This has been particularly evident in the context of serogroup C *Nme*, where immunization is associated also with a clear reduction in disease in unimmunized individuals within the vaccinated population (8, 9). Unfortunately, no capsule-based vaccine exists for serogroup B *Nme* due to similarities between the serogroup B capsular polysaccharides and human self-antigens (10). This limitation prompted the development of protein-based vaccines to target serogroup B *Nme*, including 4CMenB (11). As with conjugate vaccines, 4CMenB was approved based on SBA elicited; at the time, it was unknown whether immunization would impact nasal carriage since there remains no correlate of protection against mucosal colonization (10, 11). Mouse-based infection studies (12) and early clinical studies (11, 13, 14) following vaccine implementation now suggest that, while 4CMenB immunization protects against invasive disease, it is unlikely to impact nasal colonization on a large scale. There remains a need for improved vaccines to target group B *Nme* carriage and, ideally, a single vaccine with greater cross protection across serogroups.

While historical vaccine design tended to focus on the induction of anti-bacterial antibodies, more recent experience with upper respiratory bacterial pathogens including *Streptococcus pneumoniae* and *Bordetella pertussis* have revealed an important role for CD4^+^ T cells in mediating protection against nasal colonization. Vaccine-elicited protection against nasal colonization by *S. pneumoniae* depends on antigen-specific IL-17-producing CD4^+^ T cells that recruit neutrophils to the nose following infection (15, 16). This mucosal protection can occur in the absence of antibodies, however protection against lung infection relies on both IL-17 and antibodies (17, 18). Vaccine elicited protection against *B. pertussis* nasal colonization relies on both antigen specific TH1 and TH17 cells being present (19). Given the importance of T cells in protection against these bacterial pathogens, we sought to explore whether immunity against *Nme* nasal colonization may rely on a cellular response rather than strictly being a humoral response that confers protection against invasive disease.

A major barrier to understanding *Nme* infections has been the human-restricted nature of this pathogen. *Nme* does not naturally colonize the respiratory mucosa of any model organism. The ability of *Nme* to colonize the nasal passages of transgenic mice expressing human CEACAM1 (hCEACAM1), which is a receptor for the meningococcal Opa protein adhesins, provides an opportunity to explore meningococcal infection and immunity (20). In particular, the observation that the level of protection against nasal carriage of *N. meningitidis* in these mice reflects that observed in humans for both the serogroup C conjugate and 4CMenB vaccines (20–22) supports the utility of this experimental model.

Herein, we characterized the immune processes required to confer protection against meningococcal nasal carriage in the CEACAM1-humanized mouse model. We establish that mice must possess affinity-matured antibodies to become protected, but T cells were not required at the time of infection. Despite the central role of complement for immunity against invasive *Nme* disease, mucosal protection occurs in the absence of complement. However, immunity is strictly dependent on the presence of neutrophils at the time of infection, implicating a critical role for opsonophagocytosis on the respiratory mucosal surface. These findings advocate for the consideration of nasal immunoglobulin that promotes opsonophagocytosis as a correlate of protection against meningococcal carriage, providing both a quantifiable determinant to monitor population-level susceptibility and a target for future vaccine development.

## RESULTS

### Immunization strategies to induce protection against nasal colonization

Naïve, hCEACAM1-expressing transgenic mice are highly susceptible to nasal colonization by *Nme* (20). To establish protection against nasal colonization by *Nme*, we first compared the ability of different exposures to confer immunity. For these studies, whole bacterial cells were used as an immunogen, and mice were immunized on day 0 and 21, and challenged on day 35 (Figure 1A). Protection was evaluated based on the number of bacteria recovered from the nose 24 hours after nasal infection (on day 36).

**Figure 1.**
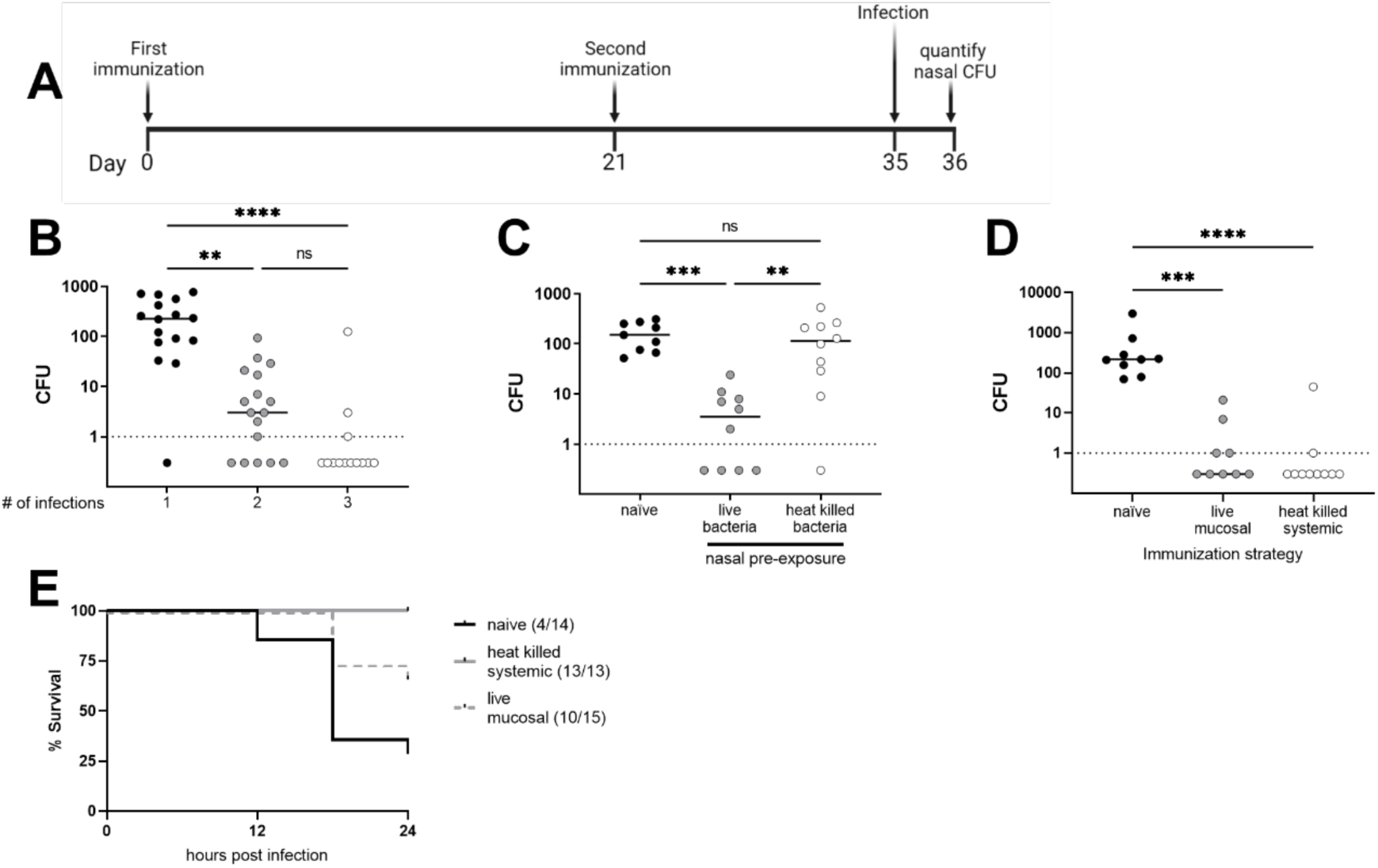
Strategies to elicit immunity against *N. meningitidis* nasal colonization. (A) An overview of the general immunization scheme used throughout this manuscript, wherein hCEACAM1 expressing mice were inoculated with live or heat-killed bacteria (as indicated) on days 0 and 21, and challenged with a live infection on day 35. (B) Nasal bacterial burdens recovered from mice 24 hours after each infection outlined in (A). Bacterial burdens are reported as colony forming units (CFU) per mouse, with each circle reflecting an individual mouse. Bacterial burdens are compared between naïve mice (black circles), mice infected two times (grey circles) and mice infected three times (white circles). (C) Mice were nasally exposed to live (grey circles) or heat-killed (white circles) *Nme* on day 0 and 21 prior to nasal challenge on day 35. Bacterial burdens were enumerated at 24 hours post infection as nasal CFU and compared to those recovered from naïve mice (black circles). (D) Mice were immunized either by live nasal infection (grey circles) or via intraperitoneal injection of heat-killed bacteria (white circles) on day 0 and 21, prior to nasal challenge on day 35. Nasal bacterial burdens were recovered at 24 hours post infection and were compared to those recovered from naïve mice (black circles). Each graph represents two independent experiments pooled, with a total of 9 to 16 mice per treatment. Bars denote group median with each point reflecting a single mouse. Statistical significance was determined by Kruskal-Wallis test with multiple comparisons. * p<0.05, ** p<0.01, *** p<0.001, **** p<0.0001. (E) Mice were immunized either by live nasal infection (grey dashed line) or heat-killed bacteria intraperitoneally (grey solid line) on day 0 and day 21, and were then challenged with a lethal dose of *Nme* on day 35 via intraperitoneal injection. Survival was compared to that of naïve mice (black solid line).

We have previously observed that nasal infection of FvB mice with *Nme* confers protection against subsequent nasal colonization (20). Here, we observed that naïve hCEACAM1-expressing C57BL/6 mice were highly susceptible to infection, with high bacterial burdens recovered from the nose at 24 hours post infection. Mice that were nasally infected once previously had reduced susceptibility to colonization upon re-exposure (second infection), evidenced by a lower nasal bacterial burden at 24 hours post-infection (Figure 1B). Most mice infected twice previously displayed near complete resistance to infection upon a third nasal challenge (Figure 1B). Since two nasal infections reproducibly conferred protection against a third nasal challenge, we used this infection scheme as a model of mucosal immunity reflecting that induced by natural colonization in humans.

When mice were intranasally exposed to heat-killed (instead of live) bacteria twice, there was no protection against colonization, indicating that a live bacterial infection is necessary for mucosal induction of immunity against nasal colonization (Figure 1C). In contrast, mice immunized intraperitoneally with heat-killed bacteria were protected from colonization following subsequent nasal challenge (Figure 1D). Intraperitoneal immunization of heat-killed bacteria was therefore used as a model of systemic immunization.

Given that any vaccine targeting *Nme* must be capable of preventing invasive disease in addition to nasal colonization, we first tested whether mucosal and systemic immunization could prevent mortality in a model of *Nme* sepsis. As expected, systemic immunization robustly protected mice against invasive challenge (Figure 1E). Repeat nasal infection was sufficient to protect the majority of immunized mice (10/15) against invasive disease but failed to elicit the same level of protection observed in mice immunized systemically (Figure 1E). This less robust protection afforded by mucosal immunization could be due to lower humoral antibody and/or SBA titers in these mice, which will be addressed below.

Therefore, when using whole bacterial cells as an immunizing agent following the immunization timeline outlined in Figure 1A, protection against *Nme* nasal colonization can be induced mucosally via two episodes of nasal carriage with live (but not heat-killed) bacteria, or parenterally via two systemic injections with heat-killed bacteria. Both methods of immunization elicited protection against invasive disease, though mucosal immunization was only protective against a low dose invasive challenge. Given the differential effects of live versus killed bacteria, and the varying levels of systemic protection achieved via nasal or systemic immunization, we sought to exploit these models to determine what factors conferred protection against nasal colonization.

### Immunization induces the production of *Nme*-specific ASC B cells and antibodies

Due to the rare and sporadic nature of invasive meningococcal disease, meningococcal vaccines have been approved based upon their ability to elicit *Nme*-specific antibodies that trigger complement-dependent bacterial killing (3). We quantified *Nme-*specific antibody responses in immunized mice to look at localization and quantity of antibody. Both mucosal and systemic immunization elicited *Nme*-specific antibodies, although the immunization strategies differed in the type and localization of antibody induced (Figure 2). Following mucosal immunization with live bacteria, high levels of *Nme*-specific IgA were detected in nasal swabs (Figure 2Aii) while no appreciable *Nme*-specific antibody response was detected in serum (Figure 2Ai). Conversely, systemic immunization elicited high levels of *Nme*-specific IgG, but not IgA, in serum (Figure 2Bi). Systemic immunization appeared to elicit low levels of *Nme*-specific IgG and IgA in nasal samples, however IgG levels did not reach statistical significance (Figure 2Bii).

**Figure 2.**
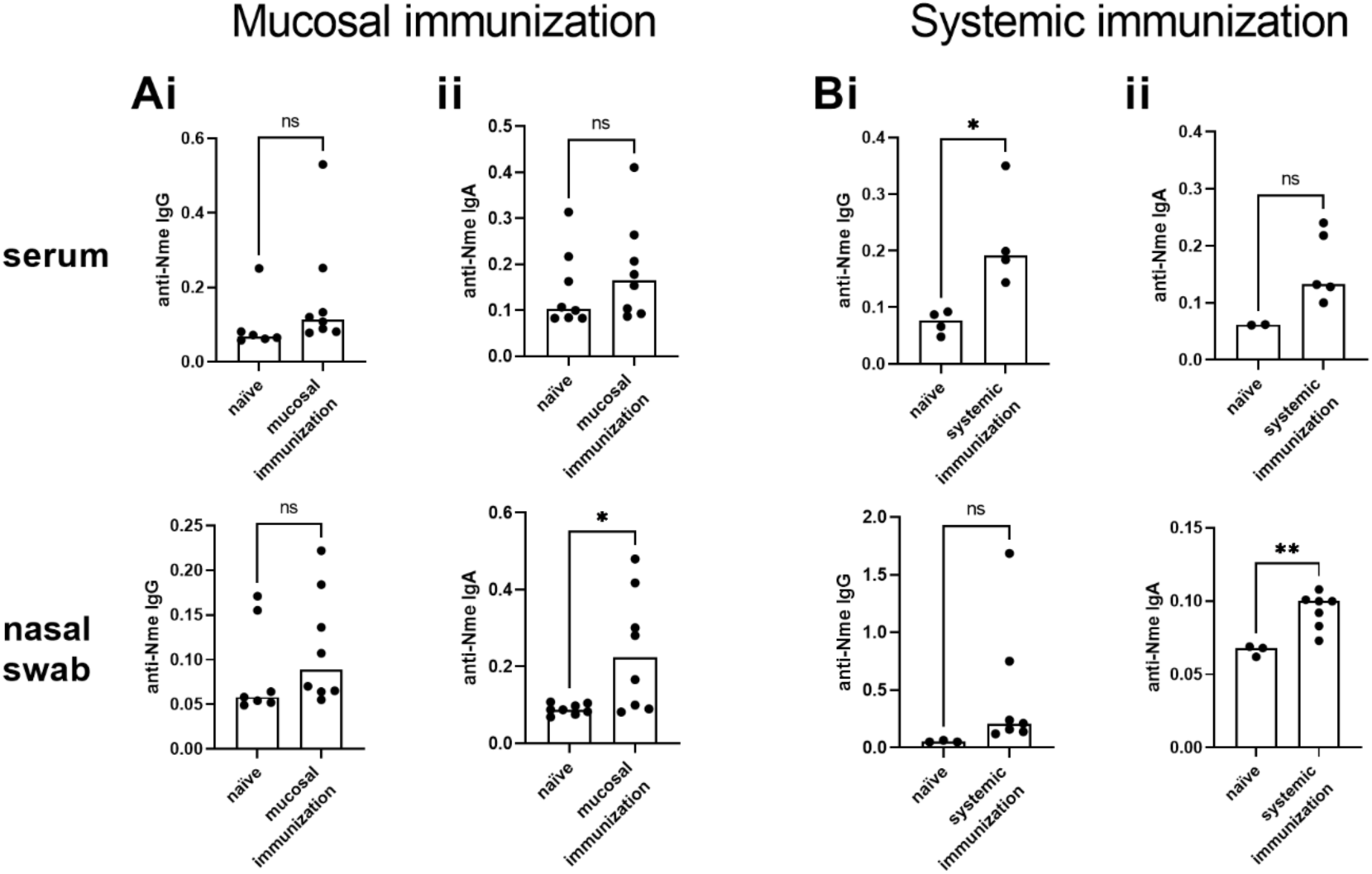
Immunization is associated with *N. meningitidis* specific antibody production. hCEACAM1 expressing mice were immunized on day 0 and 21 and were euthanized on day 35 without a terminal nasal infection. Mice were immunized via live nasal infection (A, mucosal immunization) or intraperitoneal injection with heat-killed bacteria (B, systemic immunization). *Nme* specific IgG (i) and IgA (ii) were quantified through whole bacteria ELISA and are reported as OD600 values. Bars denote group mean, with each data point reflecting a single mouse. 2-8 mice were used per group. Statistical significance was determined by unpaired t-test, * p<0.05, ns = not significant.

Mouse IgG can be characterized as one of four subclasses, IgG1, 2a or 2c (depending on the allele expressed by the mouse line), 2b, and 3 (23, 24). IgG subclasses exhibit different binding affinities to Fc receptors and, consequently, have different capacities to interact with immune cells (25). Furthermore, experimental data suggests that IgG subclasses differ in their ability to elicit killing of *Nme* (23). We therefore measured IgG subclasses elicited by each immunization strategy. Following mucosal infection, *Nme-*specific IgG2b was detected within serum and nasal swab (Figure 3A), yet no other IgG subclasses specific to *Nme* were detected in either compartment. Systemic immunization induced production of *Nme*-specific IgG2b and IgG3 (Figure 3B). These data demonstrate that both immunization strategies elicit production of *Nme* specific IgG2b, while systemic immunization also elicits production of *Nme* specific IgG3.

**Figure 3.**
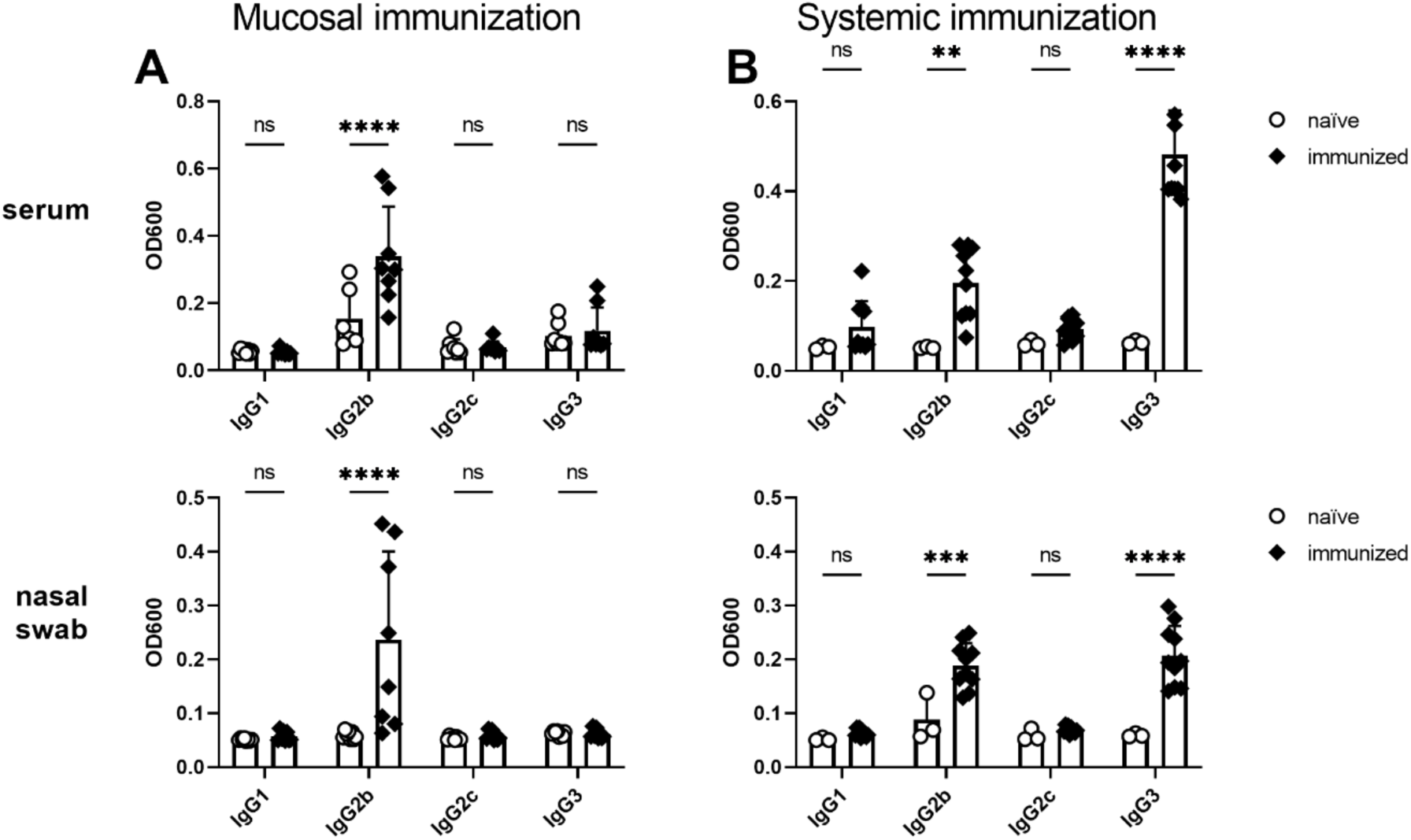
Characterization of meningococcal specific IgG subclasses following immunization. hCEACAM1 expressing mice were immunized by (A) live mucosal infection or (B) systemically with heat-killed bacteria on day 0 and 21. All mice were euthanized on day 35 without a terminal nasal infection. Serum and nasal tissue samples were collected, and meningococcal specific antibodies were quantified through whole bacterial ELISA and are reported as OD600 values. Antibodies were compared between immunized (black diamonds) and naïve (white circles) mice. Bars reflect group mean, with each point representing one mouse. 3-8 mice were used per group. Statistical significance was determined by 2-way Anova with Sidak’s multiple comparisons. * p<0.05, ** p<0.01, *** p<0.001, **** p<0.0001, ns = not significant

Given that *Nme-*specific antibodies were produced in response to immunization, but antibody isotype and localization differed depending on immunization strategy, we questioned where antibody production was taking place. To this end, we quantified IgA- and IgG-producing antibody secreting cells (ASCs) in the bone marrow (BM), cervical lymph nodes (CLN) and nasal-associated lymphoid tissue (NALT) of immunized mice. Following mucosal exposure, ASC producing *Nme-*specific IgA were observed in the CLN and NALT, and IgG-producing ASCs were observed in the CLN. No IgG-producing ASCs were observed in the NALT after mucosal infection, and no *Nme-*specific ASCs were observed in the BM of nasally infected mice (Figure 4). Systemic immunization led to high numbers of *Nme-*specific IgG-producing ASC’s in the BM, but no *Nme-*specific ASCs were apparent in the CLN or NALT (Figure 4). Interestingly, there were very few detectable IgA-producing ASC’s following systemic immunization, and these were only apparent in the BM. These data clearly demonstrate that, while both immunization strategies induce the production of *Nme* ASC’s, their location and isotype expressed is dependent on the type of immunization. Notably, these data suggest that local production of *Nme*-specific antibody (in NALT and CLN) is not required to prevent nasal colonization because mice immunized systemically are protected from colonization (Figure 1) despite only having *Nme* specific ASC in BM (Figure 4).

**Figure 4.**
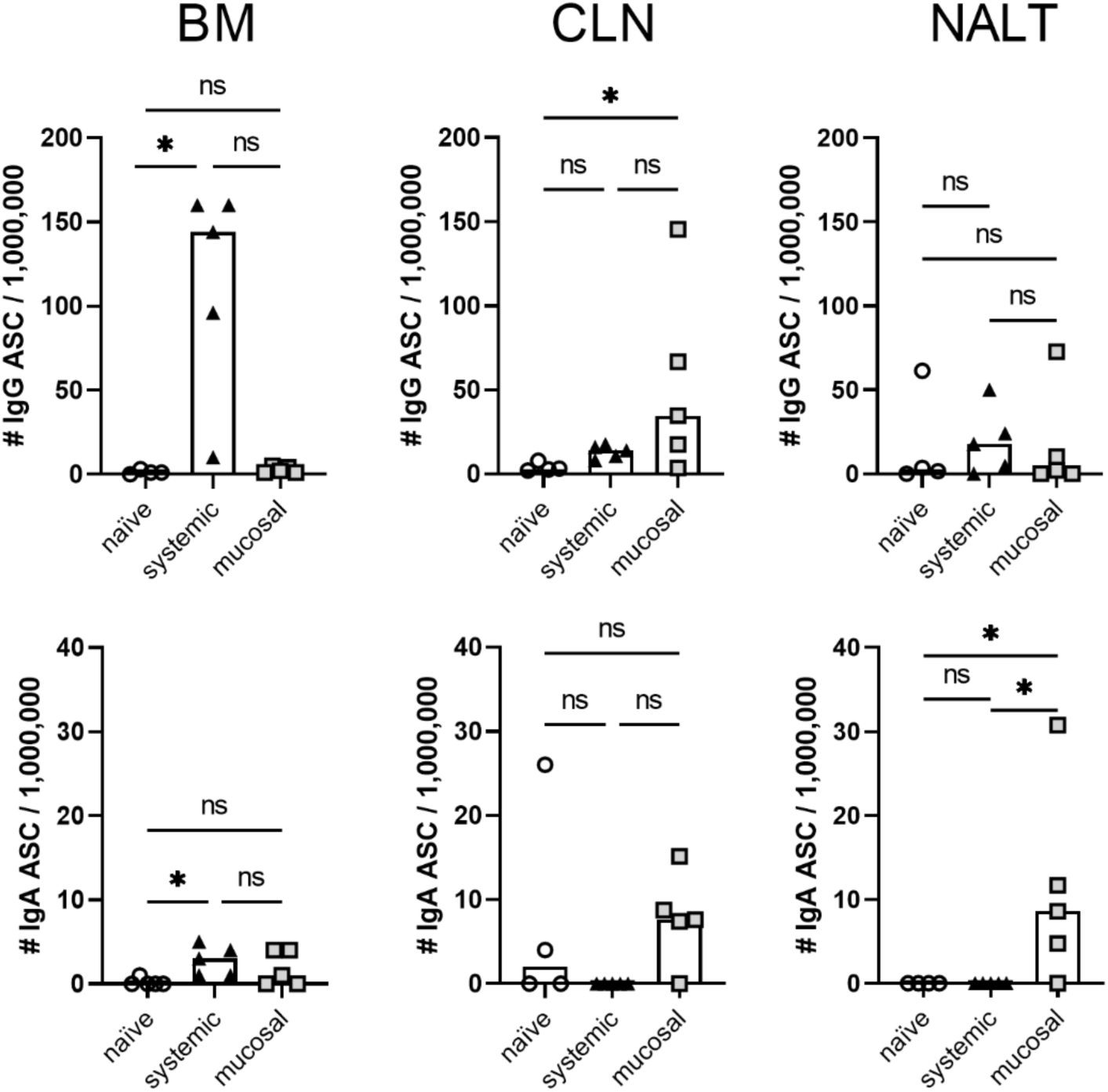
Characterization of *N. meningitidis* specific Antibody Secreting Cell (ASC) localization following immunization. hCEACAM1 expressing mice were immunized on day 0 and 21 by systemic injection with heat-killed bacteria (black triangles) or mucosal infection with live bacteria (grey squares). Naïve (white circles) and immunized mice were nasally infected on day 35 and all mice were euthanized at 24 hours post infection. Cells producing *Nme* specific IgG and IgA were enumerated in bone marrow (BM), cervical lymph nodes (CLN) and nasal associated lymphoid tissue (NALT) via two-colour ELISPOT. ASC counts are reported as a frequency of 1,000,000 cells, with symbols reflecting a single mouse and bars denoting group median. 5 mice per group. Statistical significance was determined by Kruskal-Wallis test with multiple comparisons, * p<0.05; ns = not significant.

### Protection against nasal colonization requires B cells

To address whether the production of *Nme*-specific antibodies is essential for protection against nasal colonization, we utilized mouse models lacking functional B cell responses. To this end, we immunized human CEACAM1-expressing JH knockout mice (hCEACAM1^+/-^JH^-/-^), which lack B cells and all antibody isotypes, and AID-knockout mice (hCEACAM1^+/-^AID^-/-^), which cannot undergo somatic hypermutation or class switch recombination and thus have B cells that produce non-affinity matured IgM but no other antibody isotypes (26, 27). Repeated nasal infection with *Nme* in the JH^-/-^ mice failed to elicit protection against nasal colonization, evidenced by the significantly greater nasal bacterial burden in immunized JH^-/-^ relative to B cell-producing JH^+/-^, and the lack of significant difference in immunized JH^-/-^ compared to naïve JH^-/-^ mice (Figure 5A). AID^-/-^ mice also remained susceptible to nasal colonization following mucosal exposure, while AID^+/-^ controls did become immune (Figure 5B). Systemic immunization of JH^-/-^ (Figure 5C) and AID^-/-^ (Figure 5D) also failed to induce immunity against nasal colonization. These data establish that B cells with the capacity for somatic hypermutation are an absolute requirement for protection against nasal colonization. Given that B cells are required for protection against nasal colonization, we reasoned that mice genetically deficient in T cells would be unable to develop protection against colonization (28). However, given the requirement for bacterial specific CD4^+^ T cells at the time of nasal challenge for protection against *S. pneumoniae* nasal colonization (15), we were curious to test whether a T cells were similarly involved at the time of *Nme* infection. To this end, T cell competent mice were intranasally colonized twice with live bacteria and then, prior to nasal challenge, were depleted of either CD4+ or CD8+ T cells. T cell depletion was confirmed by flow cytometry (Supplemental figure 1). T cell depletion was not uniform, however the degree of depletion did not correlate with degree of protection against colonization. T cell depletion had no impact on protection, indicating that these T cell subsets are not required for protection at the time of infection (Figure 5E). These findings reveal that, unlike *S. pneumoniae*, protection against *Nme* nasal colonization requires B cells but that CD4^+^ T cells do not obviously contribute at the time of nasal challenge.

**Figure 5.**
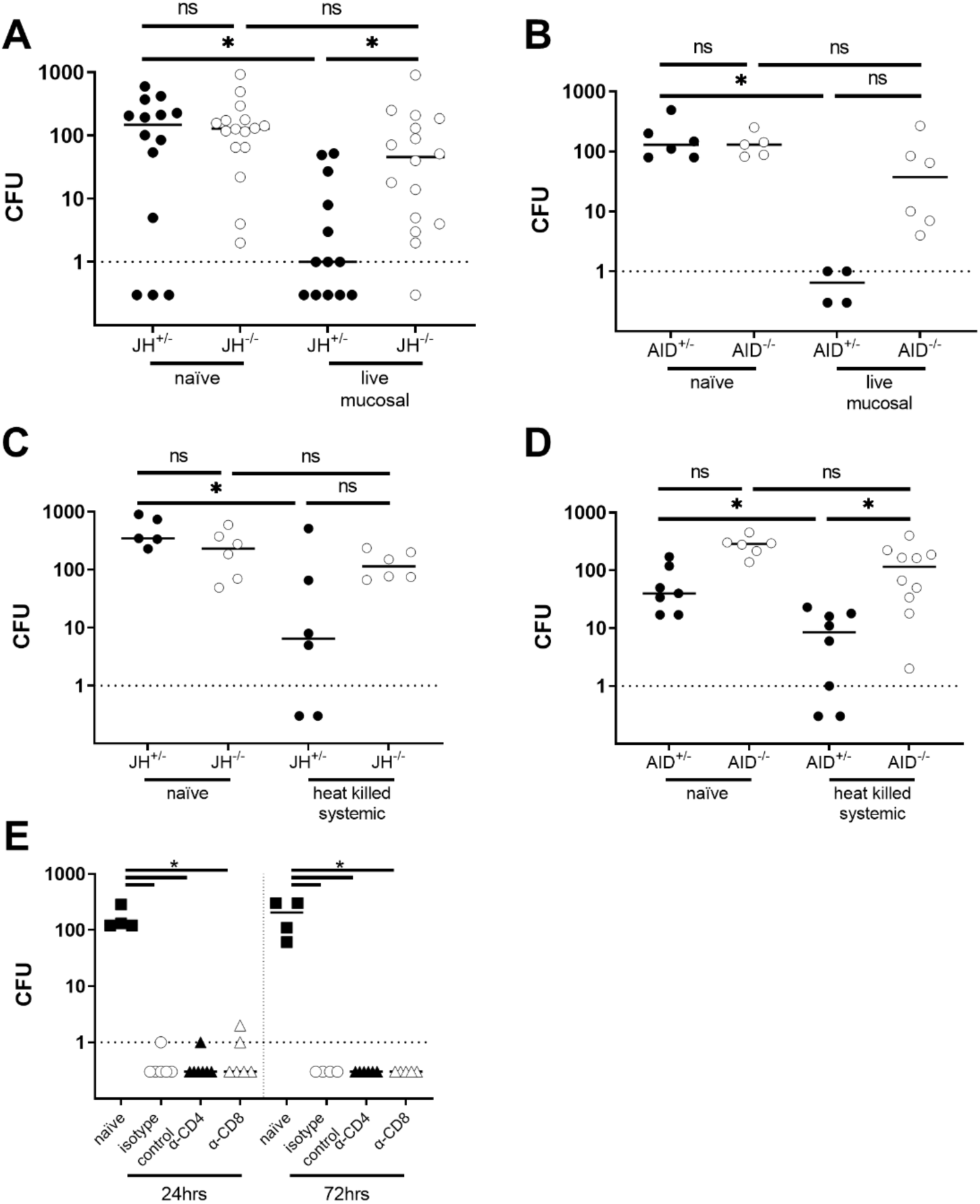
Protection against meningococcal colonization is reliant on B cells. hCEACAM1-expressing mice were immunized via mucosal infection with live bacteria (A, B and E) or via systemic injection with heat-killed bacteria (C and D) on day 0 and 21, and all mice were nasally infected on day 35. Nasal bacterial burdens were enumerated 24 hours post infection CFU and are plotted with each data point reflecting one mouse and lines denoting group medians. (A and C) Impact of immunization in B cell deficient (hCEACAM1^+/-^JH^-/-^) mice was compared to B cell competent littermates (hCEACAM1^+/-^JH^+/-^) following (A) mucosal or (C) systemic immunization. (B and D) Impact of immunization in AID deficient (hCEACAM1^+/-^AID^-/-^) mice was compared to AID competent littermates (hCEACAM1^+/-^AID^+/-^) following (B) mucosal or (D) systemic immunization. (E) The role of T cells in protection against *Nme* nasal colonization was evaluated in hCEACAM1^+/-^ mice immunized via mucosal infection. Immunized mice were depleted of T cells by injection of α-CD4 (black triangles), α-CD8 (white triangles) or isotype control antibody on day 32 and 34, prior to nasal challenge on day 35. Bacterial burdens were recovered from T cell depleted mice at 24 and 72 hours post nasal challenge. Statistical significance was determined by Kruskal-Wallis test with multiple comparisons, * p<0.05, CFU = colony forming units, ns = not significant

### The complement system is not required for protection against nasal colonization

One mechanism through which antibodies clear bacterial infections is through recruitment of complement proteins to the bacterial surface. Complement deposition subsequently triggers bacterial killing, either through the formation of the membrane attack complex (MAC), leading to cell lysis, or by opsonizing the bacteria to activate phagocytic killing (29). Serum bactericidal activity (SBA) is a quantitative measure of antibody-mediated complement lysis of bacteria. Given that SBA is used as a functional correlate of protection against invasive meningococcal disease, and vaccines against *Nme* are required to elicit an SBA response, we questioned whether SBA developed through our different immunizing approaches (3, 30). Mucosal colonization and systemic immunization both elicit a potent anti-*Nme* SBA titer (Figure 6A). However, the SBA titer induced by mucosal immunization was lower than that induced by systemic immunization, which likely explains why mucosal immunization was less protective against invasive *Nme* infection (Figure 1E).

**Figure 6.**
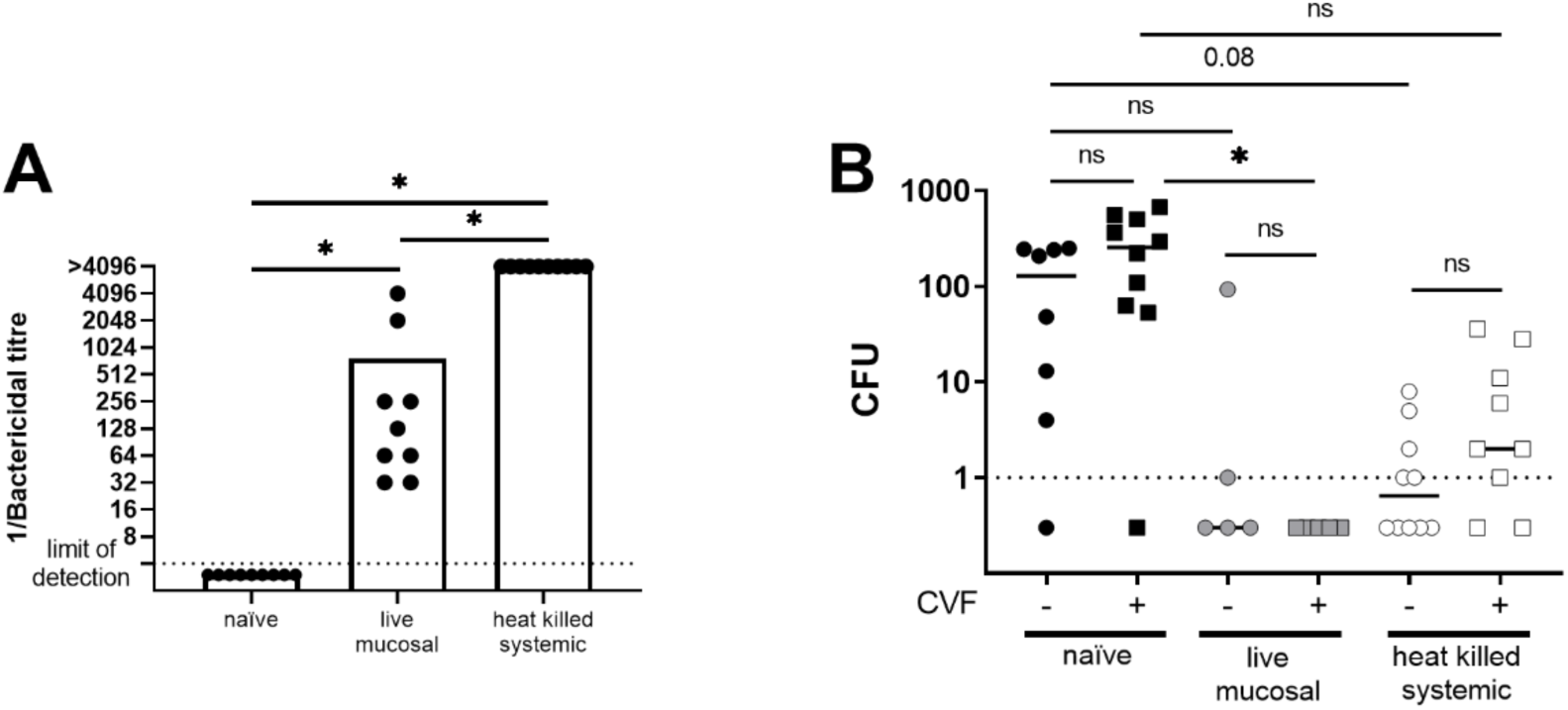
Protection against *N. meningitidis* nasal colonization can occur in the absence of complement. hCEACAM1 expressing mice were immunized via mucosal infection with live bacteria or via systemic injection with heat-killed bacteria on day 0 and 21. (A) Serum from immunized and naïve mice was collected on day 35 without nasal challenge. 1/Bactericidal titre is plotted as the highest dilution of serum which resulted in a 50% or more reduction in viable bacteria compared to no serum controls. Bars denote group means, each symbol reflects an individual mouse. The lowest dilution of serum tested was 1:4 (limit of detection), serum was then sequentially diluted two-fold to a maximum dilution of 1:4096. Samples that failed to kill 50% of bacteria at any dilution of serum are plotted below the limit of detection while samples that killed 50% or more bacteria at a dilution of 1:4096 are plotted as >4096. Bars denote group means with symbols reflecting a single mouse. (B) Complement was depleted by CVF injection 24 hours prior to infection. Nasal bacterial burdens were quantified as colony forming units (CFU) at 24 hours post infection. Bacterial burdens were compared between complement depleted (CVF + mice; squares) and complement competent (CVF – mice; circles) mice. Bacterial burdens were compared between naïve (black) mice or those immunized through mucosal infection (grey) or systemic injection with heat-killed bacteria (white). Lines reflect group median and each symbol reflects the bacterial burden recovered from a single mouse. Colonization data is represented as CFU per mouse, with lines denoting median bacterial burden. Statistical significance was determined by Kruskal-Wallis test with multiple comparisons, * p<0.05. CFU = colony forming units, ns = not significant

Given that immunization led to development of *Nme-*specific antibodies that effectively promote complement-dependent bacterial killing, we next tested whether complement was required for protection by injecting mice with cobra venom factor (CVF) to deplete complement C3 prior to infection, thus preventing both C3-mediated opsonophagocytosis and MAC-dependent lysis. Complement depletion was confirmed by Western blot (Supplemental figure 2A). As we have previously observed, complement depletion in naïve mice had no significant impact on bacterial burden in comparison to untreated mice (Figure 6B and (20)). Notably, immunized mice were effectively protected from nasal colonization in the absence of complement, evidenced by the lower bacterial burden in the immunized but complement-depleted mice compared to complement-depleted naïve mice (Figure 6B). This indicates that complement is not required for protection against nasal colonization, despite its clear importance in protection against invasive disease. Interestingly, while complement depletion in systemically-immunized mice did not abolish protection against nasal colonization, there was an observable increase in the proportion of mice with recoverable bacteria in the nose relative to the untreated (normal complement) immunized mice (Figure 6B). These data may suggest that protection afforded by systemic immunization optimally relies on the complement system, but that effective protection can occur in the absence of the C3-triggered complement system. This is consistent with the fact that IgG2b, the most prevalent IgG subclass detected in immunized mice (Figure 3), does not activate complement mediated killing of bacteria.

### Immune effector expression following nasal infection suggests neutrophil recruitment

To characterize what type of immune response is associated with bacterial clearance following infection, we quantified immune effector expression in nasal lavages and serum by ELISA. Expression of immune effectors was unchanged in serum following infection (data not shown), indicating that the inflammation remains localized in response to nasal infection. Mucosal and systemic immunization strategies both led to a marked increase in myeloperoxidase (MPO) and elastase 2 (ELA2) in nasal tissues at 6 hours post-infection (Figure 7A and B). By 16 hours post-infection, naïve mice had comparably high MPO and ELA2 (Figure 7C and D), though this appears to be due to a combination of a slow increase in naïve mice and a decline in peak response in the immunized animals at this later time point. MPO and ELA2 are both produced by neutrophils, and the high expression of these effectors suggest neutrophil influx following nasal infection (31).

**Figure 7.**
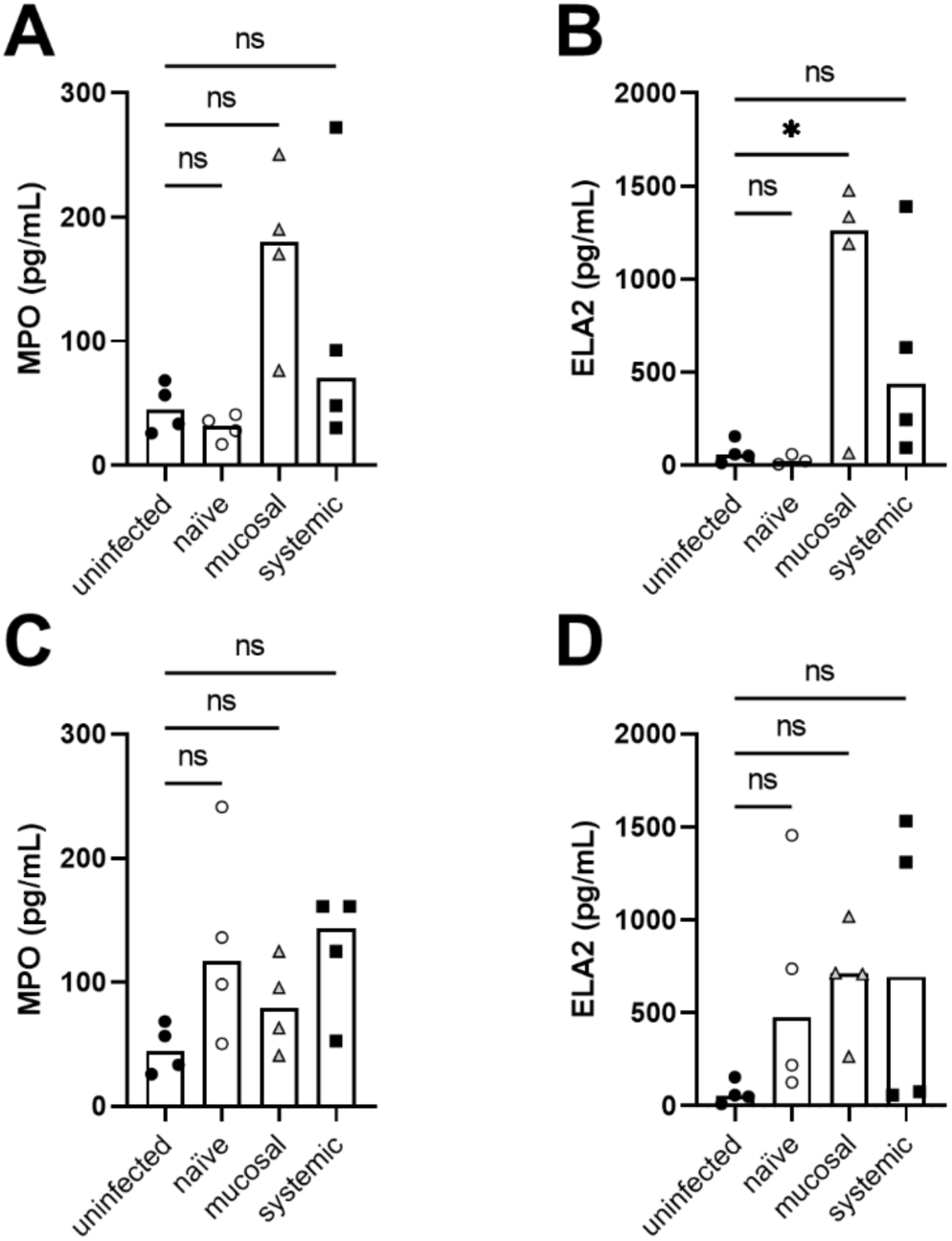
Quantification of inflammatory enzymes following *N. meningitidis* infection. hCEACAM1 expressing mice were immunized via live mucosal infection (grey triangles) or via systemic injection with heat-killed bacteria (black squares) on day 0 and day 21, and were nasally challenged on day 35 with naïve controls (white circles). Mice were euthanized at 6 (A and B) and 16 (C and D) hours post infection and nasal lavages were collected. Myeloperoxidase (MPO) and Elastase2 (ELA2) levels were quantified via ELISA and compared between infected mice and uninfected (black circles) controls). Shown here are representative graphs from 1 of 3 independent replicates, with 4 mice per treatment group, bars denoting median and each symbol representing a single mouse. 2 uninfected mice collected at each timepoint and pooled to give 4 uninfected mice. Statistical significance was determined via one-way Anova with Sidak’s multiple comparisons. * p<0.05, ns = not significant.

### Neutrophils are required for protection against nasal colonization

Given the elevated expression of neutrophil effector molecules in the nasal passages of immunized mice following *Nme* infection, and that neutrophils express Fc receptors that bind to IgG2b to trigger phagocytosis of invading microbes (25), we were curious how much antibody-mediated phagocytosis contributed to protection against *Nme* mucosal colonization. CEACAM-humanized mice were therefore immunologically-depleted of neutrophils prior to infection, with the effectiveness of depletion confirmed by flow cytometry (Supplemental figure 2B). This resulted in an increased bacterial burden in all mice (Figure 8A and B). The increased bacterial burden in naïve mice in the absence of neutrophils reveals an important role for neutrophils in controlling early stages of *Nme* nasal colonization, even in the absence of specific antibodies. Neutrophil depletion in immunized mice resulted in a loss of protection against nasal colonization, evidenced by the increase in bacterial burden relative to undepleted immunized mice, revealing an essential role for neutrophils in conferring the adaptive immune protection against nasal colonization. Importantly, immunized mice that were neutrophil depleted had a lower bacterial burden than did naïve neutrophil-depleted mice, suggesting that some level of protection persisted in these immunized mice in the absence of neutrophils. However, these data support a critical role for neutrophils in mediating innate and adaptive protection against nasal colonization.

**Figure 8.**
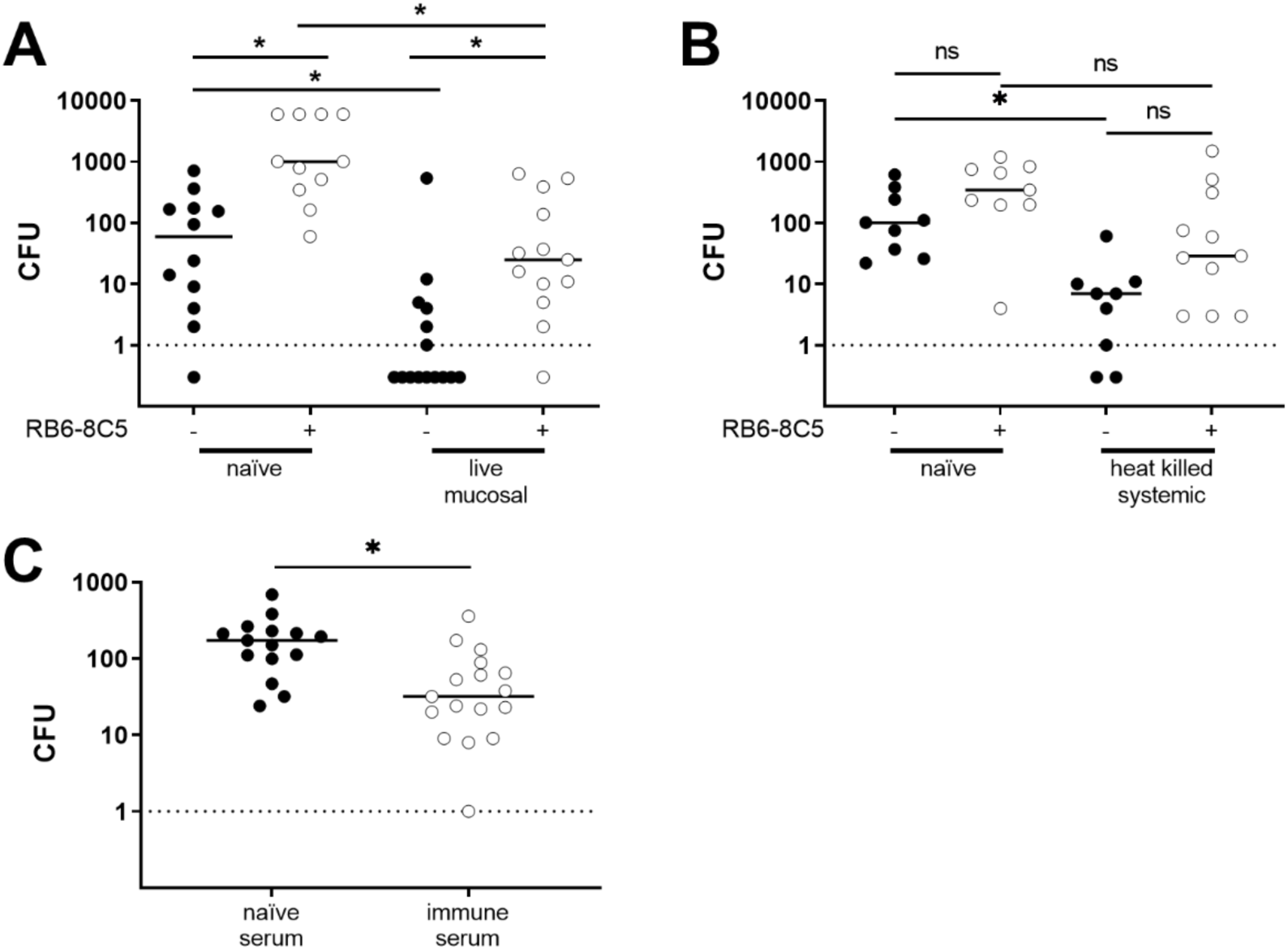
Protection against *N. meningitidis* nasal colonization is reliant on neutrophils. Mice were immunized via live mucosal infection (A) or systemic injection of heat-killed bacteria (B) on day 0 and 21 and were then nasally infected on day 35. Neutrophils were depleted via intraperitoneal injection of RB6-8C5 on day 32 and 34. Nasal bacterial burdens were compared between naïve and immunized mice, and neutrophil depleted (white circles) and neutrophil competent (black circles) mice at 24 hours post infection. Statistical significance was determined via Kruskal-Wallis with Dunn’s multiple comparisons test. (C) Serum for passive transfer was generated through systemic immunization, wherein wildtype mice were intraperitoneally injected with heat-killed bacteria on day 0 and 21 and serum was collected on day 35. Naïve, hCEACAM1-expressing mice were intraperitoneally injected with 100 µL of serum from naïve (black circles) or systemically immunized (white circles) 24 hours prior to infection. These mice were then nasally infected and nasal bacterial burdens were enumerated 24 hours post infection. Statistical significance was determined by Mann-Whitman. Lines reflect group median and each symbol reflects the bacterial burden recovered from a single mouse. * p<0.05, ns = not significant.

### Serum transfer from immune mice confers protection against nasal colonization

We next tested whether antibodies were the only adaptive immune component required to confer protection against nasal colonization. To this end, heat-inactivated serum from systemically immunized or naïve mice was injected into naïve mice 24 hours prior to *Nme* nasal infection. Following infection, mice that received serum from systemically immunized mice had a significantly lower nasal bacterial burden than mice who received naïve serum (Figure 8C). While the protection afforded was not as complete as that conferred by active immunization, this clearly establishes that antibody can itself afford protection.

## DISCUSSION

The remarkable success of polysaccharide conjugate vaccines in reducing the burden of invasive meningococcal disease is reliant on both the ability of these vaccines to prevent nasal colonization, leading to a community-protective herd immunity, and to protect vaccinated individuals against systemic infection (4, 9). More recently developed vaccines use protein antigens to elicit protection against serogroup B meningococci, which cannot be targeted by polysaccharide-based vaccines (10, 32). Unfortunately, the lack of any known correlate of protection against nasal colonization has made it impossible to deliberately focus on nasal colonization during vaccine development, despite this being a necessary first step to disease. Post-implementation studies have revealed that these protein-based vaccines can prevent *Nme* invasive disease in vaccinated individuals, but do not have a major impact on nasal colonization (11, 13, 21, 33). Here, we used CEACAM1-humanized mice to reveal what immune processes are required to protect nasal colonization, with hopes that a greater understanding of these processes will facilitate improved vaccine design against *Nme* and other respiratory pathogens.

Protection against nasal colonization elicited by conjugate vaccines is reliant on polysaccharide-specific antibodies, a fact evident because T cells cannot recognize polysaccharide antigens and most protein carriers used for the conjugate are irrelevant to the vaccine-targeted pathogens (28). The very high density of these repeating sugars on the bacterial surface provides an optimal opsonophagocytic target; based upon our results herein, this undoubtedly contributes to their ability to provide sterilizing immunity. In contrast, protein vaccines tend to target lower density epitopes but can elicit development of pathogen-specific B and T cells, both of which can directly contribute to protection. This distinction is most clearly supported by the finding that immunization with protein antigens can confer protection against nasal colonization by *Streptococcus pneumoniae* even in the absence of antibodies (19, 34). Here we observed that protection against meningococcal nasal colonization required functional, affinity-matured and/or class-switched B cells and an antibody response, since neither AID^-/-^ not JH^-/-^ mice were protected against nasal colonization following mucosal or systemic immunization. Given that *S. pneumoniae* and *Nme* colonize the same mucosal surfaces, it is remarkable that these differences exist (15, 16, 34).

Passive transfer of immune serum into naïve mice resulted in a reduced bacterial burden relative to mice that received naïve serum, indicating that antibodies alone can confer protection against *Nme* nasal colonization (Figure 8C). The protection afforded by passive transfer was not as effective as that afforded by active immunization; all mice that received the immune sera became colonized, while active immunization resulted in most mice having no detectable *Nme*. The lower level of protection afforded by passive transfer may be due to technical limitations, such as the amount of antisera transferred and/or its ability to be transported to the mucosal surface in the time frame of these experiments, but it may also indicate that the presence of *Nme-*specific adaptive immune cells (T or B) do contribute to optimal control of the infection.

Immunity could be attained by either nasal infection or intraperitoneal injection of killed bacteria. While both elicited IgG and IgA, mucosal immunization resulted in local antibody production, with *Nme*-specific ASCs in the cervical lymph nodes and NALT of immunized mice, while systemic immunization resulted in distal antibody production, with *Nme-*specific ASCs confined to the bone marrow. Both approaches led to the emergence of *Nme*-specific antibodies in the serum and nares prior to nasal challenge.

Antibodies from both immunization strategies promoted complement-dependent killing of *Nme* and, accordingly, both conferred protection against invasive infection. Serum bactericidal activity is an effective correlate of protection against invasive *Nme* disease, so meningococcal vaccines have been approved for use in humans based upon their ability to elicit an SBA response (10, 11, 35). However, some protein vaccines that elicit a significant bactericidal response against *Nme* do not impact nasal colonization (11, 13, 21, 33, 35, 36), which indicates that SBA should not be considered a correlate of protection against nasal colonization.

Beyond SBAs simply being an inadequate predictor of protection against colonization, complement depletion in immunized mice did not result in a loss of protection against nasal colonization, suggesting that mucosal immunity against *Nme* can occur in the absence of complement-dependent processes. However, given that complement can act as an opsonin, and our data suggests that opsonophagocytosis is required for effective protection against nasal colonization, it seems likely that complement deposition on the bacterial surface may still facilitate clearance of the meningococci. Consistent with this, previous studies have demonstrated that complement enhances phagocytic killing of *Nme* in blood (29, 37). This may also explain why complement depletion in the parenterally immunized mice led to a modest, but not statistically significant, increase in susceptibility to nasal colonization in this model.

While neutrophils are generally considered to be important for innate defense against bacterial infections on the mucosa, we here establish that they are also required for the prevention of nasal colonization by *Nme* in immunized mice. Neutrophils are also an important effector in the vaccine-mediated protection against nasal colonization by *S. pneumoniae*. With the pneumococci, acapsular whole cell vaccine-elicited T_h_17 cells secrete IL-17 in response to nasal infection, which subsequently induces rapid recruitment of neutrophils to the nasal passages (16, 38). A loss of either CD4^+^ T cells or IL-17 resulted in a loss of protection against nasal colonization (16, 34), while the loss of B cells (& antibodies) did not (34, 39). With meningococcal infection, we also observed that the neutrophil response was more rapid in immunized mice, and the neutrophils were required for the protection afforded by both mucosal infection and systemic immunization. Since antibodies were also required for immunity, this suggests that the antibody-mediated opsonization enhances phagocytic clearance of the meningococci by neutrophils, as illustrated in Figure 9. Since depletion of CD4^+^ T cells did not impact protection against nasal colonization by *Nme*, this clearly differentiates protection from that observed in the *S. pneumoniae* model. It is enticing to consider that the meningococci’s ability to avoid complement deposition on its surface, including via host factor H binding (40), explains the enhanced requirement for antibodies as an opsonin within the mucosal tissues.

**Figure 9.**
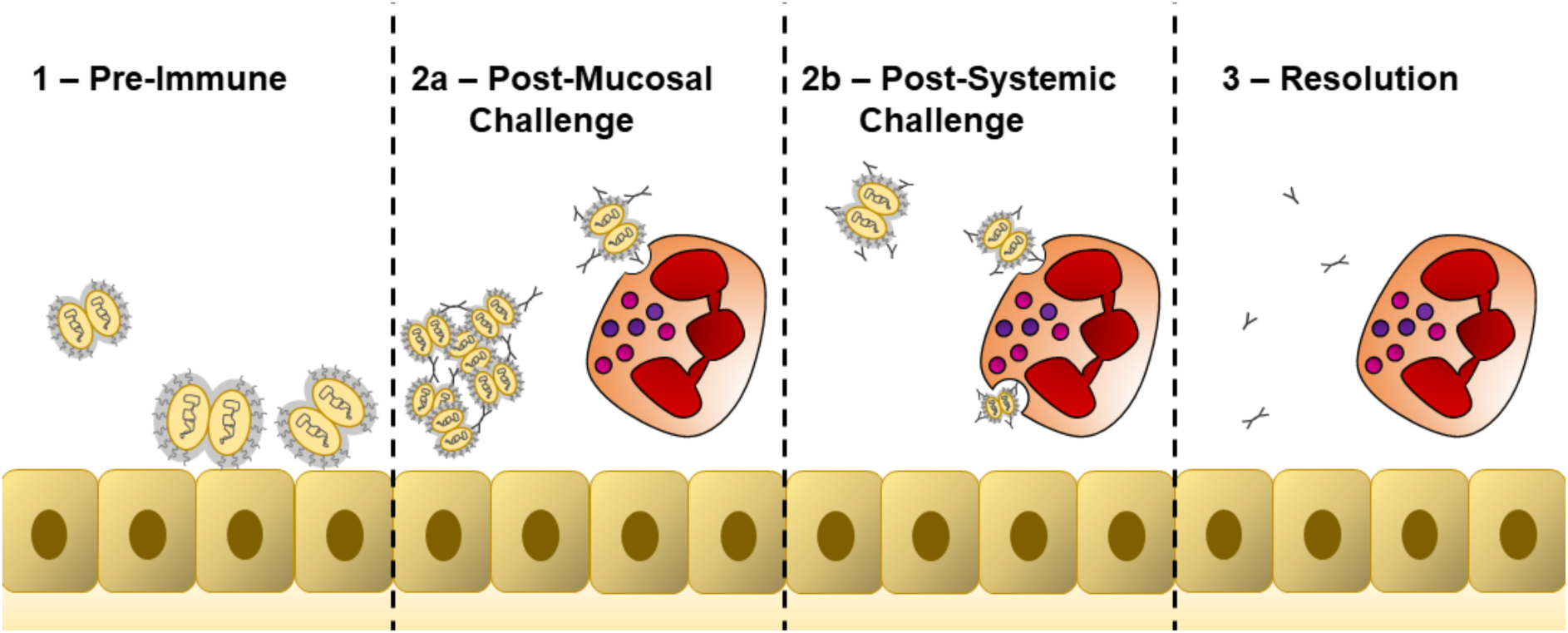
Overview of vaccine-elicited protection against meningococcal nasal colonization. (1) Pre-immune mice possess no meningococcal antibodies and *Nme* effectively colonizes the nasal epithelium. (2a) Mice immunized via live infection (mucosal) possess nasal *Nme* specific IgA and IgG. Following nasal challenge, *Nme* specific IgA can lead to bacterial clumping. *Nme* specific IgG can interact with Fc receptors on neutrophils to facilitate neutrophil dependent bacterial clearance. (2b) Mice immunized via heat-killed bacteria injected intraperitoneally (systemic) possess *Nme* specific IgG. Following nasal infection, antibody-coated bacteria can efficiently be phagocytosed by neutrophils, resulting in neutrophil-dependent bacterial clearance. (3) Immunized mice rapidly clear nasal *Nme* infections.

For both meningococcal vaccine development and monitoring of population-level immunity, it is critical to consider correlates that reflect the potential for sterilizing immune protection. Our data suggests that effective clearance of *Nme* nasal colonization relies on antibody-mediated opsonophagocytosis by neutrophils. Complement deposition on the bacterial surface might enhance this process, but it is not sufficient to mediate clearance in the absence of antibodies or neutrophils. These findings align with the prescribed use of meningococcal vaccines in individuals who carry complement deficiency, a practice known to be effective when using polysaccharide conjugate vaccines against the meningococci (29, 41). They also help to explain why some protein-based, parenterally-administered meningococcal vaccines that promote a significant serum bactericidal response and protect against invasive meningococcal disease are not effective against nasopharyngeal carriage. This work also supports the premise that mucosal protection afforded by natural infection can be monitored by the presence of *Nme-*specific nasal IgA, whereas parenterally-administered vaccines instead confer protection via a systemically-produced IgG response. If these effects reflect that in humans, then the titer and ratio of *Nme-*specific immunoglobulin isotypes will reveal an individual’s susceptibility to infection and differentiate between natural and vaccine-induced immunity.

## MATERIALS AND METHODS

### Bacterial growth conditions

*Nme* serogroup C strain 90/18311 was used for all studies. *Nme* was routinely grown on GC agar (Becton Dickinson) supplemented with IsoVitalex (Becton Dickinson) and, when recovering bacteria from mice, VCNT inhibitor (Becton Dickinson), incubated at 37°C with 5% CO_2_. For growth in broth, BHI (Becton Dickinson) was supplemented with 1% IsoVitalex and incubated at 37°C with agitation.

### Mouse infection studies

All animal experimental procedures were approved by the Animal Ethics Review Committee of the University of Toronto, which is subject to the ethical and legal requirements under the province of Ontario’s Animals for Research Act and the federal Council on Animal Care (CCAC).

FvB mice that express human CEACAM1 (hCEACAM1 (42)) were backcrossed for 12 generations onto the C57BL/6 background, allowing interbreeding with immune deficient mutants available in this line. These transgenic animals were used for all infection studies. JH^-/-^ mice were crossed with hCEACAM1^+^ mice to generate JH^-/-^ hCEACAM1^+^ mice. Heterozygous JH^+/-^ hCEACAM1^+^ littermates were used as B cell competent controls for comparison. AID^-/-^ mice were crossed with hCEACAM1^+^ to generate AID^-/-^ hCEACAM1^+^ and AID^+/-^ hCEACAM1^+^ mice. All mice were bred inhouse and littermates were used as controls for all studies. Unless otherwise stated, female and male mice were used in equal numbers.

Nasal and invasive infections were performed as previously described (20). For nasal infections, awake mice were nasally inoculated with 1 x 10^5^ CFU of bacteria in 10 µL PBS with 1 μM MgCl_2_ and 0.5 μM CaCl_2_ (PBS/Mg/Ca). Mice were euthanized 24 hours post infection for bacterial enumeration, or at 6- and 16-hours post-infection for cytokine analysis. Nasal bacterial burden was quantified via nasal swab. Nasal swabs were collected in 500 µL sterile PBS/Mg/Ca, which were centrifuged at 3000 rpm, with resulting pellets resuspended and plated to enumerate bacterial burden while supernatant was saved for further analysis. Serum was collected via cardiac puncture and nasal lavage was collected post-sacrifice by retrograde lavage with 200 µL sterile PBS through the trachea and out of the nose. Serum, lavage and nasal swab supernatant were collected on ice and stored at -20°C until further analysis.

For invasive studies, male hCEACAM1 mice were intraperitoneally challenged with 1 x 10^4^ *Nme* and supplemented with a separate IP injection of 2 mg iron dextran in sterile PBS to provide an iron source to support bacterial replication. Mice were monitored for clinical symptoms at least every 12 hours and humanely euthanized when a pre-determined endpoint was met.

### Immunization studies

Mice were immunized on day 1 and 21, via injection of heat-killed bacteria or by intranasal administration of heat-killed or live bacteria, followed by nasal infection on day 35 (Figure 1A). To generate heat-killed bacteria for immunization purposes, an overnight culture of *Nme* was resuspended in BHI supplemented with 1% IsoVitalex and incubated at 37°C with agitation until the culture reached log phase, defined by OD_600_ (∼0.6). Cultures were then killed at 60°C for 40 minutes after which heat-killed bacteria was aliquoted and stored at - 80°C until use. For immunizations with heat-killed bacteria, mice were intraperitoneally injected with 1 x 10^7^ CFU in 200 µL PBS/Mg/Ca or nasally inoculated with 1 x 10^5^ CFU in 10 µL PBS, following the protocol for live nasal infection as indicated above. Intranasal immunization with live bacteria was performed by repeated intranasal administration as described above.

### *N. meningitidis*-specific whole bacterial ELISA

*Nme*-specific antibodies were quantified as described elsewhere (43). Briefly, heat-inactivated *Nme* was diluted to an OD_600_=0.2, and used to coat 384-well flat bottom immune non-sterile plates (Thermo Scientific). 20 µL of undiluted sample (mouse serum or nasal swab supernatant) was applied to each well. The quantity of *Nme-*specific antibody was measured using the following detection antibodies: Alkaline Phosphatase (AP) conjugated-goat-anti-mouse IgG Fc(γ) (Jackson 115-055-008), AP-goat-anti–mouse IgM (Jackson 115-055-020), AP-goat-anti-mouse IgA (Abcam ab97232), AP-goat-anti-mouse IgG1 (Jackson 115-055-205), AP-goat-anti-mouse IgG2a (Jackson 115-055-206), AP-goat-anti-mouse IgG2b AP (Jackson 115-055-207), AP-goat-anti-mouse IgG2c AP (Jackson 115-055-208) and AP-goat-anti-mouse IgG3 AP (Jackson 115-055-209).

## ELISPOT

Membrane plates (0.45 µm Hydrophobic High Protein Binding Immobilon-P Membrane, Millipore) were coated with approximately 10^5^ CFU/mL heat-killed *N. meningitidis* and placed at 4°C overnight. The plates were blocked with 10% FBS RPMI for at least 2 hours at 37°C. Single cell suspensions derived from bone marrow, cervical lymph nodes and nasal-associated lymphoid tissue were loaded onto the plate at serial 2-fold dilutions in 10% FBS RPMI, starting with 1 million cells, and incubated overnight at 37°C. The next day cells were removed, and wells were washed 5 times with PBS containing 0.1% TWEEN-20. HRP-conjugated IgA (Southern Biotech, 2714213) and AP-conjugated IgG (Southern Biotech, 2692322) detection antibodies were subsequently added for 2 hours at 37°C.

Plates were then washed 3 times with PBS containing 0.1% TWEEN-20, and a further 3 times with PBS. Plates were developed in the dark until spots were visible using AEC Peroxidase (for HRP-conjugated antibodies, Vector Laboratories) and Vector Blue (for AP-conjugated antibodies, Vector Laboratories) substrates. After development plates were washed with distilled water and were left to dry overnight prior to spot enumeration. Spots were counted based on original cell dilution and were converted to number of spots (ASCs) per million.

### Detection of neutrophil associated enzymes via ELISA

At 6- and 16-hours post-infection nasal lavage was collected on ice and stored at -20°C until analysis. Mouse neutrophil Elastase (ELA2) and mouse myeloperoxidase (MPO) were quantified in undiluted samples by ELISA (R&D Systems).

### Serum bactericidal activity

Serum bactericidal activity was measured using a modified version of a previously published protocol (44). Serum was heat inactivated at 56°C for 30 minutes and diluted two-fold in Hanks Buffered Salt Solution with CaCl_2_ and MgCl_2_ (Gibco), starting at a dilution of 1:4. Overnight cultures of *Nme* were re-streaked onto GC agar with IsoVitalex for 4 hours. Bacteria and baby rabbit complement were added in equal volume to diluted serum. Reactions were incubated at 37°C for 1 hour, after which bacteria were plated to measure viability. Serum dilutions resulting in at least 50% decrease in viable bacteria compared to no serum controls were considered bactericidal. Serum bactericidal titer was plotted as the inverse of the final dilution of serum to give a 50% decrease in bacterial viability.

### In vivo depletion of T cells, complement and neutrophils

For T cell depletions, monoclonal antibodies were purified from hybridomas GK1.5 (anti-CD4) and 53-6.72 (anti-CD8) (ATCC). Mice were intraperitoneally injected with 200 µg of antibody in 500 µL of sterile PBS three times, on days 5, 3 and 1 prior to nasal infection, based on a previously outlined protocol (17). Neutrophils were depleted by injection of 250 µg of monoclonal anti-Gr.1 antibody (purified from RB6-8C5 hybridoma) two times, on days 3 and 1 prior to nasal infection (20). Expected depletion of immune cells was confirmed by flow cytometry.

Complement depletion was achieved through injection of 20 µg of Cobra Venom Factor (CVF) (Quidel) in 200 µL of PBS 24 hours prior to infection (20). Depletion of complement was confirmed by Western blot.

### Passive transfer

Serum for passive transfer was collected from naïve mice and from mice that were immunized with heat-killed bacteria (“immune serum”) as described above. Serum was pooled from 10 mice per treatment group and heat inactivated at 56°C for 30 minutes. Heat-inactivated serum was stored at -20°C until use. For passive transfer, mice were intraperitoneally injected with 100 µL of pooled serum 24 hours prior to infection. Nasal bacterial burdens were quantified as described above at 24 hours post infection.

### Statistical analysis

All statistical analyses were performed using GraphPad Prism version 9.1 (GraphPad Software).

## Acknowledgements

We are grateful to Carolyn Buckwalter, Jamie Fegan, Stacey Xu, Steven Ahn, Nelly Leung and Jessica Lam for their technical support and helpful discussions throughout the course of this work. We would also thank the Department of Comparative Medicine at the University of Toronto for their outstanding support of the animal work.

